# A meta-analysis of ancient and present-day Central Eurasian genome data to revise archaic hominin ancestry

**DOI:** 10.64898/2026.08.10.743976

**Authors:** Aigerim Rymbekova, Martin Kuhlwilm

## Abstract

Archaic introgression has shaped the evolutionary history of Eurasian populations, yet Central Eurasian region remains understudied despite being at the crossroads of ancient human migration. Here, we analyzed the whole-genome data of five Central Eurasian (CE) individuals from Early Bronze Age (EBA) and five present-day CE individuals to characterize the archaic introgression landscape. We estimated that archaic introgression from Neanderthal and Denisovan archaic hominins comprises approximately 2.2% of the Central Eurasian genomes. Both amount and chromosomal distribution of archaic introgression remained largely unchanged between the EBA and present-day CE individuals. Putative introgressed fragments matching the Altai Neanderthal and the Altai Denisovan were retrieved. Our results suggest that while the archaic introgression levels seemingly remained stable over the past several thousand years, larger modern CE genomes panels will be required to fully characterize the genomic landscape of archaic ancestry in the region.

## 1 Introduction

Central Eurasia is a region which experienced extensive migration from both Western and Eastern Eurasia throughout history and has been home to modern humans and the archaic hominins since at least the Late Pleistocene [Krause et al., 2007, Reich et al., 2010, Prufer and et. al, 2014, Haak et al., 2015, Damgaard et al., 2018, Narasimhan et al., 2019]. Introgression from Neandertals and Denisovans has shaped the evolutionary history of Eurasian populations with selection and phenotypic traits, including skin and hair pigmentation in present-day Europeans [Dannemann and Kelso, 2017] and high-altitude adaptation in Tibetan populations [Huerta-Sanchez et al., 2014]. While European and East Asian populations have been extensively studied, archaic introgression has not been addressed in previous studies on Central Eurasia (CE). Notably, the massive westward expansion of East Asian populations during the Mongol Empire dramatically altered the ancestry composition of CE populations, shifting the proportions of West and East Asian ancestry from approximately 75/25 to nearly 50/50 [Jeong et al., 2019, Narasimhan et al., 2019]. As East Asian populations carry introgressed variation from possibly several Denisovan-like lineages [Browning et al., 2018], the substantial East Asian ancestry influx may have introduced a unique set of archaic variation into the CE populations. Whether this ancestry shift is reflected as a detectable change in the amount or composition of archaic introgression between Early Bronze Age (EBA) and present-day CE genomes remains unclear.

Archaic ancestry represents a minor fraction of modern human genomes, hence reconstruction of the temporal landscape of archaic introgression in CE requires multiple genomes sampled at different points in time. The availability of a small set of medium to high coverage EBA and high coverage present-day CE genomes (five individuals from each time period) makes it possible to start comparing archaic ancestry across this ancestry transition. However, much larger sample sizes are needed to investigate any temporal trends. Here, we used whole-genome sequencing data from present-day [Kairov et al., 2022] and ancient individuals [Damgaard et al., 2018] (Table 1) from CE to assess the variation in the individual genomes using an archaic ancestry inference tool, admixfrog, and inferred genomic regions in a “human-typical”, “Neandertal-typical” or “Denisovan-typical” state. admixfrog was chosen over other archaic introgression detection methods because it can be applied to mapped reads of individual ancient genomes directly, without requiring genotype calls of high-coverage, phased whole-population genomes, making it well-suited to CE samples where such population-scale reference data are not available. This allowed us to estimate the proportion of introgressed material from each archaic source inherited from archaic hominins during Bronze Age as well as in the present.

**Table 1:** The overview of individuals analyzed in this study.

| Individual ID | Period | Coverage | Approx. Date | Source |
| --- | --- | --- | --- | --- |
| BOT14 | EBA | 3.7x | 3371-3354 BCE | [Damgaard et al., 2018] |
| BOT15 | EBA | 2.4x | 3327-3094 BCE | [Damgaard et al., 2018] |
| BOT2016 | EBA | 13.6x | 3521-3377 BCE | [Damgaard et al., 2018] |
| EBA1 | EBA | 3.2x | 2286-2037 calBCE | [Damgaard et al., 2018] |
| EBA2 | EBA | 7.1x | 2620-2468 calBCE | [Damgaard et al., 2018] |
| KAZ_WG2 | Present-day | 33.1x | — | [Kairov et al., 2022] |
| KAZ_WG4 | Present-day | 30.49x | — | [Kairov et al., 2022] |
| KAZ_WG5 | Present-day | 27.77x | — | [Kairov et al., 2022] |
| KAZ_WG6 | Present-day | 26.71x | — | [Kairov et al., 2022] |
| KAZ_WG7 | Present-day | 28.66x | — | [Kairov et al., 2022] |

## 2 Results

The archaic introgression analysis in CE individuals revealed the genomic landscape of archaic ancestry, using admixfrog. For each individual, the genome was partitioned into non-overlapping windows and assigned an ancestry state, based on estimated posterior probabilities, allowing both the total genomic proportion and the chromosomal distribution of putative archaic fragments to be compared across individuals.

The total amount of archaic introgression estimated by admixfrog was approximately 2.2% of the genome, including both Neanderthal and Denisovan regions, across both EBA and present-day individuals (Fig. 1 and Fig. 2). The total length and chromosomal distribution of putative archaic fragments were highly consistent across individuals within each group and remained largely similar between EBA and present-day groups overall. Given that, a single representative karyogram from each group is shown (Fig. 3).

**Figure 1:**
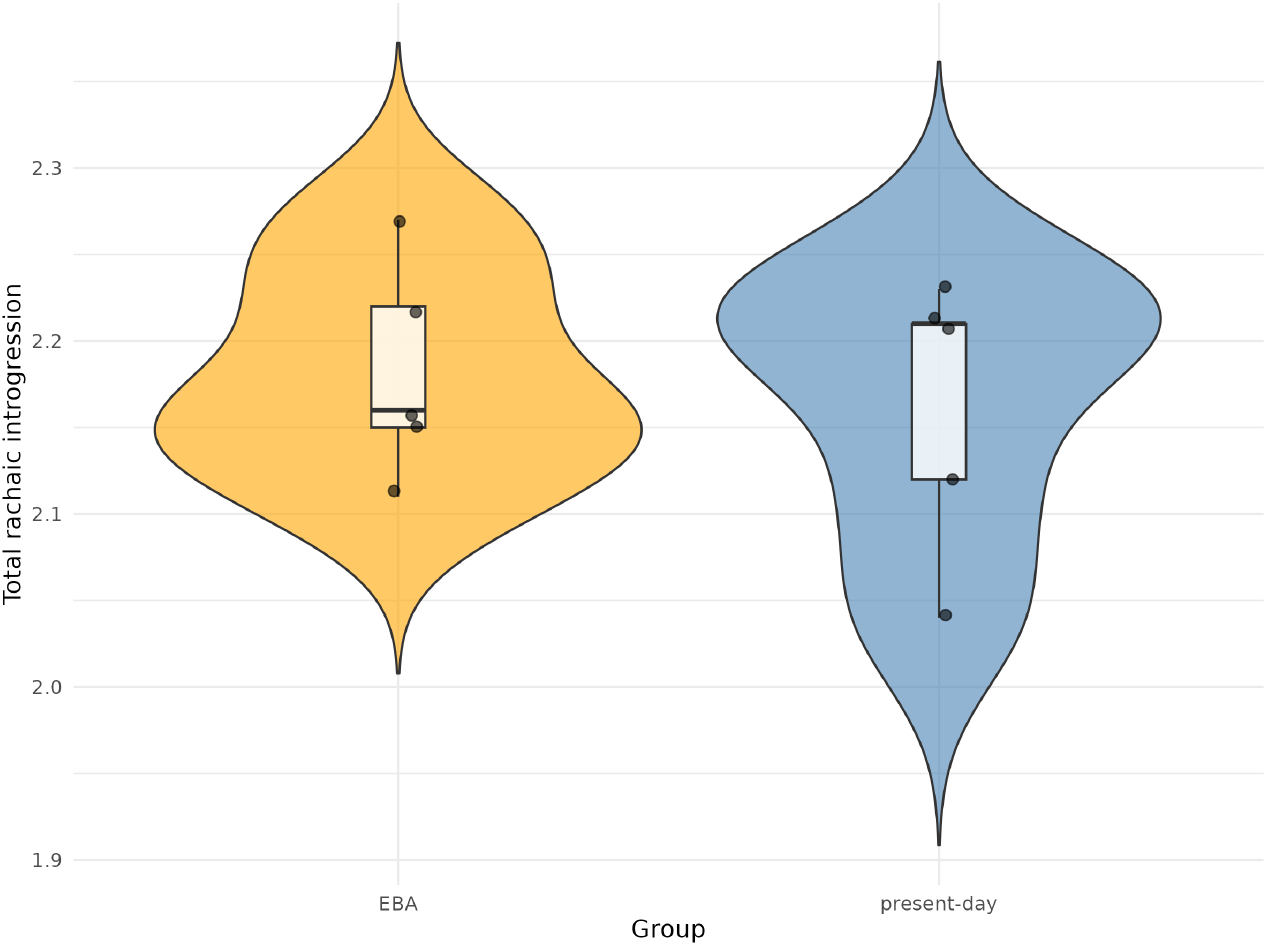
Distribution of summed Neanderthal and Denisovan ancestry proportions per individual, grouped by population (EBA, n=5; present-day, n=5). Violin plots show the estimated density of total archaic ancestry within each group; embedded boxplots indicate the median and interquartile range, with whiskers extending to 1.5× the IQR.

**Figure 2:**
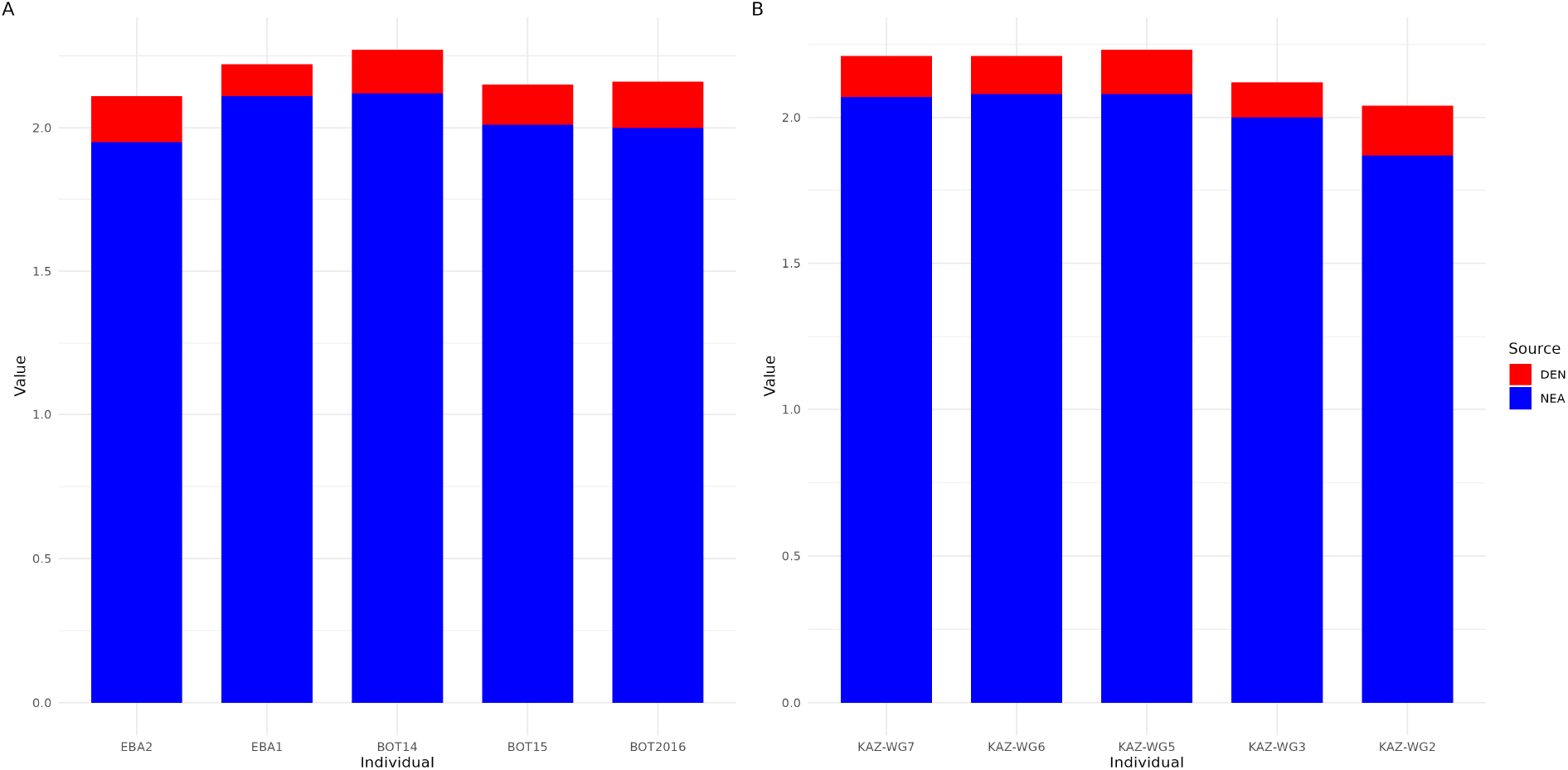
Estimated Neanderthal (NEA, blue) and Denisovan (DEN, red) ancestry proportions per individual shown as stacked bar plots. A. Early Bronze Age individuals. B. Present-day individuals. Bar height represents the total combined archaic ancestry per individual, with each segment reflecting the contribution from each archaic source.

**Figure 3:**
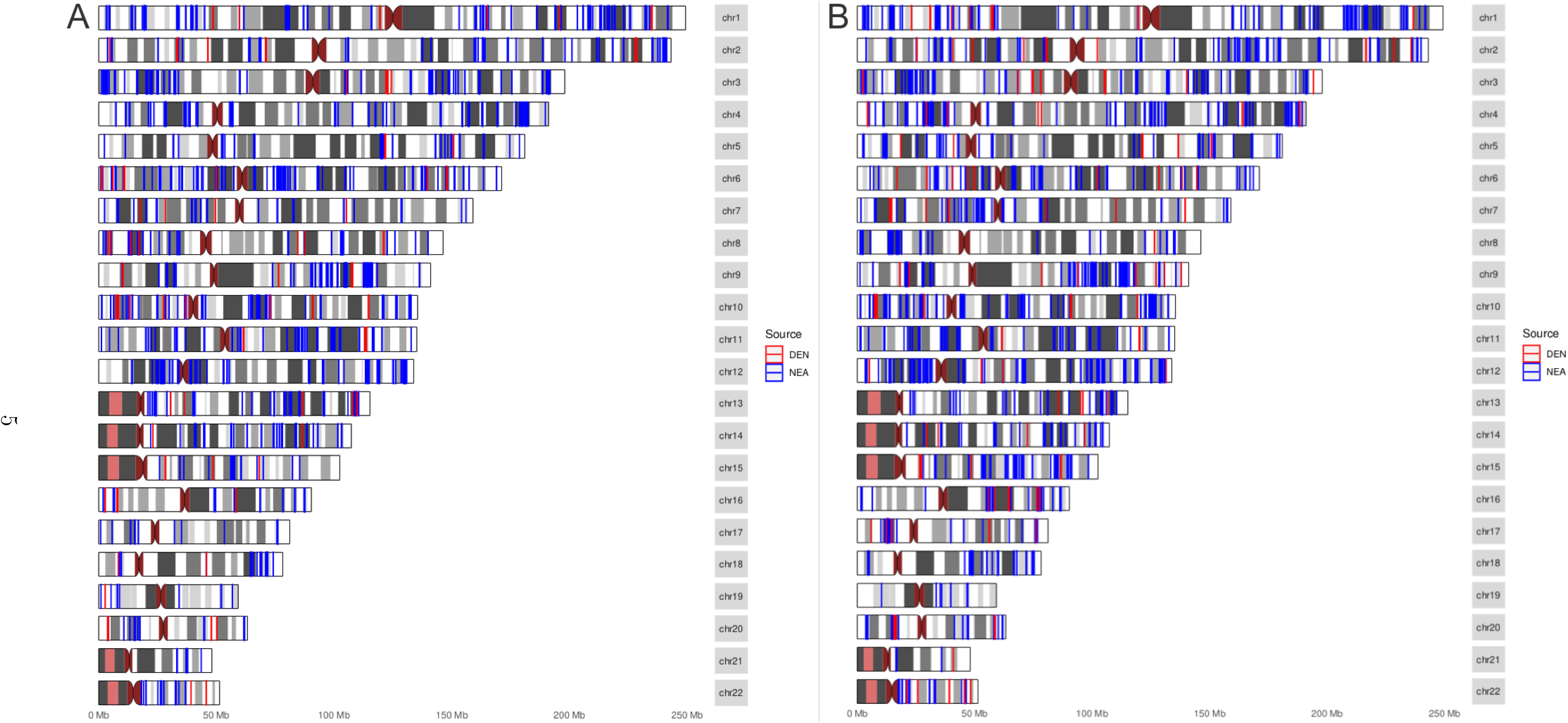
Distribution of putative archaic introgression across the genome inferred using admixfrog method, colored blue (Neanderthal) and red (Denisovan). Chromosomes are stacked along the Y-axis and chromosome length is indicated along the X-axis. A. The karyogram of an ancient Botai individual (BOT14) from Early Bronze Age. B. The karyogram of a present-day Central Eurasian individual (KAZ_WG5).

Putative archaic fragments were distributed across nearly all autosomes rather than being concentrated in a small number of chromosomal regions in all individuals examined. Neanderthal-typical fragments (blue) were consistently more numerous and in most chromosomes longer than Denisovan-typical fragments (red) across both groups. No large, chromosome-scale differences in archaic fragments placement were observed between EBA and present-day groups, and no outlier individuals were identified within each group.

## 3 Discussion

Total archaic ancestry (∼2%) estimated for CE individuals falls within the range reported in European and East Asian populations, indicating that CE genomes do not carry a distinct archaic ancestry profile despite repeated large-scale admixture events. However, the stability of archaic ancestry proportions despite of the ancestry shift may reflect several factors. First, if the East Asian ancestry introduced during the Mongol Empire-era expansion carried archaic fragments similar in magnitude to the one already present in the CE region, admixture would not be expected to substantially alter total archaic ancestry. Second, our analysis is limited by sample size of five individuals per period and by differences in sequencing coverage between ancient and modern genomes. Both constraints reduce power of detecting subtle shifts in archaic ancestry, particularly for Denisovan-typical fragments, which occur at lower frequency and typically of shorter length than Neanderthal-typical fragments. Finally, other methods for detecting introgression might be more suitable to obtain precise estimates, especially regarding Denisovan introgression, where the available Denisovan genome is a poor match to the introgressing source population Browning et al. [2018].

The latter is directly relevant to the possibility of a second, East Asian-associated Denisovan-like lineage in CE genomes. As shown in Browning et al. [2018], there were potentially multiple genetically distinct Denisovan-like introgression events in East Asian and Oceanian populations. If a comparable lineage was introduced into Central Eurasia during East Asian ancestry influx, it would unlikely be detected using the individuals present in this study. Thus, the absence of a measurable change in Denisovan-typical ancestry does not constitute evidence against the presence of a distinct Denisovan-like lineage, rather constraints of low sample size and sequencing depth used in this analysis.

## 4 Conclusion

This study represents an initial exploration of archaic ancestry in the CE region. Future studies including high-coverage CE genomes will facilitate resolving the extent and variety of archaic introgression in this understudied region. Inferring introgressed regions and connecting the variation within these regions to potential phenotypic and disease outcomes will be valuable to understand the impact of archaic admixture relative to other Eurasian populations.

## 5 Methods

### 5.1 Admixfrog analysis

The detailed scripts and necessary files are available at the corresponding GitHub repository: https://github.com/admixVIE/Eurasian-archaic-introgression.

#### 5.1.1 Central Eurasian data processing

The BAM files of ancient CE individuals mapped to the hg19 human reference genome (GRCh37) were downloaded from the European Nucleotide Archive (ENA) browser https://www.ebi.ac.uk/ena/browser/view/PRJEB26349. Raw FASTQ data of modern CE individuals were downloaded from the Sequence Read Archive (SRA) https://www.ncbi.nlm.nih.gov/bioproject/PRJNA374772. The raw reads were trimmed using Trimmomatic (version 0.39) [Bolger et al., 2014] and mapped to the hg19 human reference genome with the Burrows-Wheeler Aligner (BWA) mem algorithm (version 0.7.19-r1273) [Li and Durbin, 2009]. Polymerase chain reaction (PCR) duplicates were removed using Picard MarkDuplicates (version 3.1.1) [Institute, 2025].

#### 5.1.2 Reference data processing

The published VCF files of archaic hominins, including Denisovan [Meyer and et. al, 2012] and Neanderthal [Prufer and et. al, 2014, 2017] genomes, were obtained from https://cdna.eva.mpg.de/neandertal/Vindija/VCF/, and the corresponding mask files from https://cdna.eva.mpg.de/neandertal/Vindija/FilterBed/ were applied. The files were then filtered using an archaic admixture ascertainment BED file and subset to only genotype (GT) field using bcftools (version 1.23.1) [Danecek et al., 2021].

The individual VCF files of Mbuti Africans mapped to the hg19 human reference genome were downloaded from https://reichdata.hms.harvard.edu/pub/datasets/sgdp/. The files were merged into a single multi-individual VCF file, with missing sites filled as homozygous reference (0/0), split by chromosome.

Finally, raw FASTQ files of a chimpanzee (ERR1709961) [de Manuel et al., 2016] were obtained from https://www.ebi.ac.uk/ena/browser/view/PRJEB15086 and processed as described in [Han et al., 2025].

To process the modern human data for merging with the archaic data, archaic variant positions were first retrieved and used to construct a temporary VCF file for each autosomal chromosome, containing a single dummy individual with all genotypes set to homozygous reference. These temporary VCF files were used as a scaffold to retrieve African genotypes at archaic positions only, with missing genotypes set to homozygous reference. The dummy individual was subsequently removed, and the files were subset to retain only genotype field as well.

The chimpanzee data was processed in a similar manner prior to merging, retrieving archaic positions only using the scaffold of a temporary VCF file and filtering to retain only the genotype field.

Finally, the three datasets were merged per autosome using bcftools merge: the archaic VCFs filtered for the GT field, the modern human VCFs subset to archaic positions only and filtered for the GT field, and the chimpanzee VCF file processed in the same manner.

#### 5.1.3 Running admixfrog

Admixfrog (version 0.7.2) [Peter, 2020] analysis was performed following the documentation, available at https://github.com/BenjaminPeter/admixfrog/. The reference file was created using the *admixfrog-ref* subcommand on the merged reference VCF file. Input files were generated using the *admixfrog-bam* subcommand on the target BAM files. The analysis was run per individual using the *admixfrog* command with the reference file. The output was visualized using a custom R (version 4.3.1) script [Team, 2021].

## Data availability

This study used only previously published and publicly available data. No new sequence data were generated.

## Acknowledgments

The computational work of this study was performed using the Life Science Compute Cluster (LiSC) of the University of Vienna.

## Author contributions

A.R. performed data analysis and wrote the manuscript. M.K. supervised the work.

## Research funding

This project has been funded by the Vienna Science and Technology Fund (WWTF) [https://doi.org/10.47379/VRG20001] to M.K.

